# Synergetic impacts of turbulence and fishing reduce ocean biomass

**DOI:** 10.1101/2021.10.04.459351

**Authors:** Jody C. McKerral, Justin R. Seymour, Trish J. Lavery, Paul J. Rogers, Thomas C. Jeffries, James S. Paterson, Ben Roudnew, Charlie Huveneers, Kelly Newton, Virginie van Dongen-Vogels, Nardi P. Cribb, Karina M. Winn, Renee J. Smith, Crystal L. Beckmann, Eloise Prime, Claire M. Charlton, Maria Kleshnina, Susanna R. Grigson, Marika Takeuchi, Laurent Seuront, James G. Mitchell

## Abstract

A universal scaling relationship exists between organism abundance and body size^1,2^. Within ocean habitats this relationship deviates from that generally observed in terrestrial systems^2–4^, where marine macro-fauna display steeper size-abundance scaling than expected. This is indicative of a fundamental shift in food-web organization, yet a conclusive mechanism for this pattern has remained elusive. We demonstrate that while fishing has partially contributed to the reduced abundance of larger organisms, a larger effect comes from ocean turbulence: the energetic cost of movement within a turbulent environment induces additional biomass losses among the nekton. These results identify turbulence as a novel mechanism governing the marine size-abundance distribution, highlighting the complex interplay of biophysical forces that must be considered alongside anthropogenic impacts in processes governing marine ecosystems.

## Main Text

Across all ecosystems, a fundamental scaling relationship exists between species abundance (*A*) and body size (*W*), whereby:

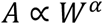

and the exponent *α* typically approximates −0.75^1^. This universal rule derives from resource acquisition as a function of body size^1^, which is a barometer for ecosystem health that simplifies interactions in complex food webs and may direct fisheries management^5^. However, within marine ecosystems, the exponent for this relationship often differs from that in terrestrial ecosystems^2^. Life-history, trophic strategies, altered productivity, and fisheries are all proposed to alter the scaling slopes of both species size-abundance distributions and individual size spectra^2–4,6^. Here, we quantify, empirically and with an independent model, how fishing and ocean turbulence cause qualitatively distinct breaks in the global marine size-abundance distribution.

For the scaling analysis we compiled size-abundance data for 2179 species, ranging from viruses to blue whales. Analyses were undertaken on a database built from primary literature and online databases (*n* = 15,146 datapoints), with secondary verification undertaken using the manually curated literature data alone (*n* = 1719) to ensure there was not systematic bias in the online sources (Methods); additionally, fits were undertaken through a balanced subsampling routine to ensure a diverse spread of species and sizes (Methods). As previously observed within individual size spectra^3^, nonlinearity was apparent in the log-transformed global size-abundance plot (Figure 1a). This coincided with a statistically verified break in the scaling value at the plankton-nekton transition of *l* ≃ 0.1 m (*l* = 0.08 m, 95% CI (0.06, 0.1)) (Methods). The marine virus to marine invertebrate slope at *α* = −0.77 is comparable to terrestrial slopes^7^. However, for organisms ≥ 0.1 m *α* was −1.9 (Figure 1a, Table 1), representing a significant negative perturbation in the slope. A shift in biomass would only translate the line downward (i.e. change the intercept via a step break), but the large slope break evidenced by these two exponent values (Figure 1) is indicative of a more fundamental alteration in the mechanistic processes shaping the species size-abundance distribution and ecosystem structure.

**Figure 1:**
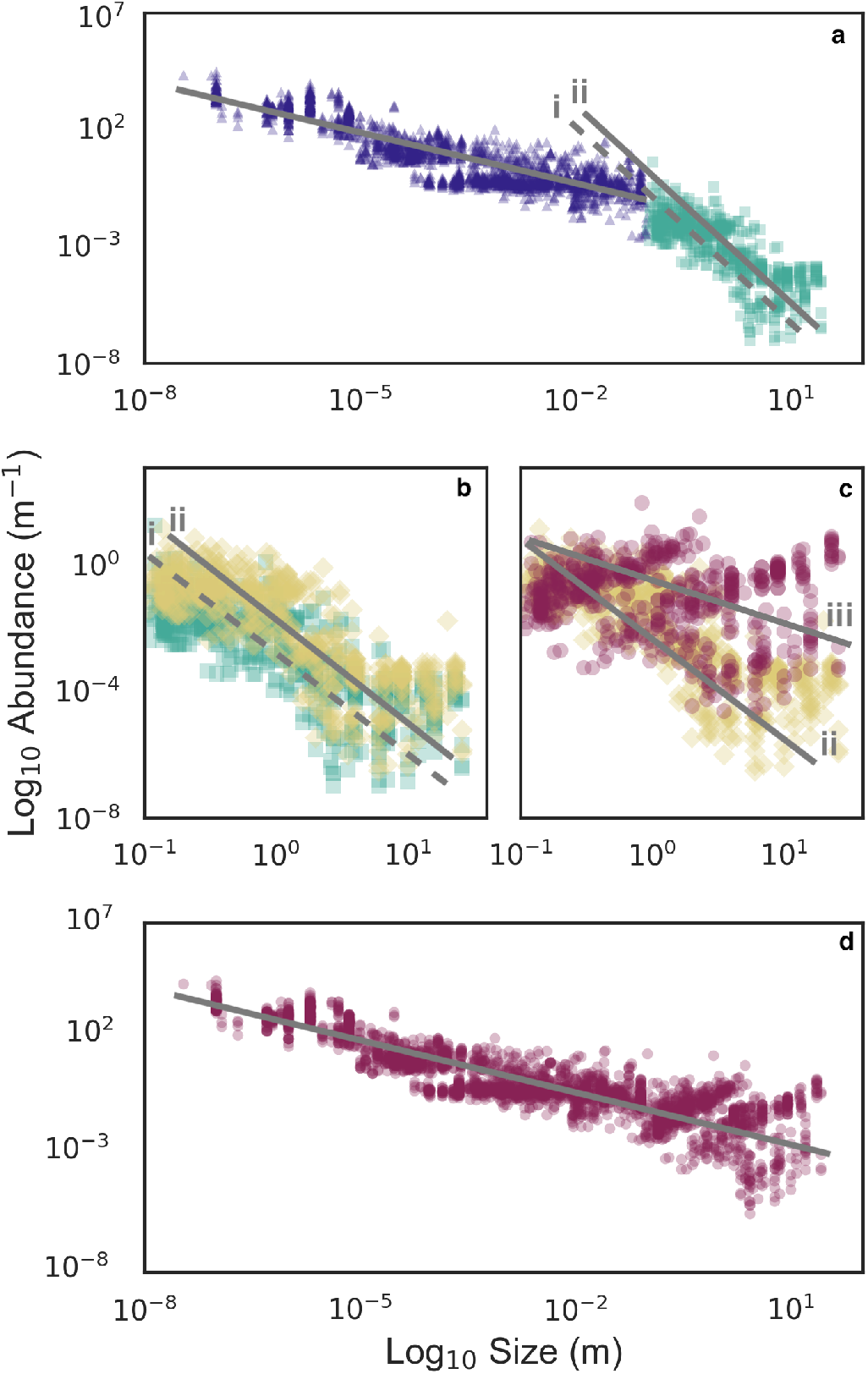
Empirical size-abundance distribution. For consistency of units, here we use length scales rather than mass/density; it is to be noted that as the transformation of both axes is the same, the relationship between size and abundance remains unchanged (Methods). **(a)** Size versus abundance for viruses to blue whales. There is a break in the scaling relationship at 0.1m. Blue triangles represent plankton ranging from viruses to zooplankton and invertebrates. Green squares are fished nekton, ranging from small fish to whales. Line (i) is the best fit to the scaling of nekton. **(b)** Line (ii) is corrected abundance for removal by fishing, with yellow diamonds showing the fishing-corrected values. **(c)** Line (iii) corrects for extra foraging due to turbulent dispersion, shown by red circles. (i) to (ii) is predominantly a vertical translation and (ii) to (iii) is a slope correction. After both corrections all points fall along a line with a slope of −0.73 **(d)**.

**Table 1.**
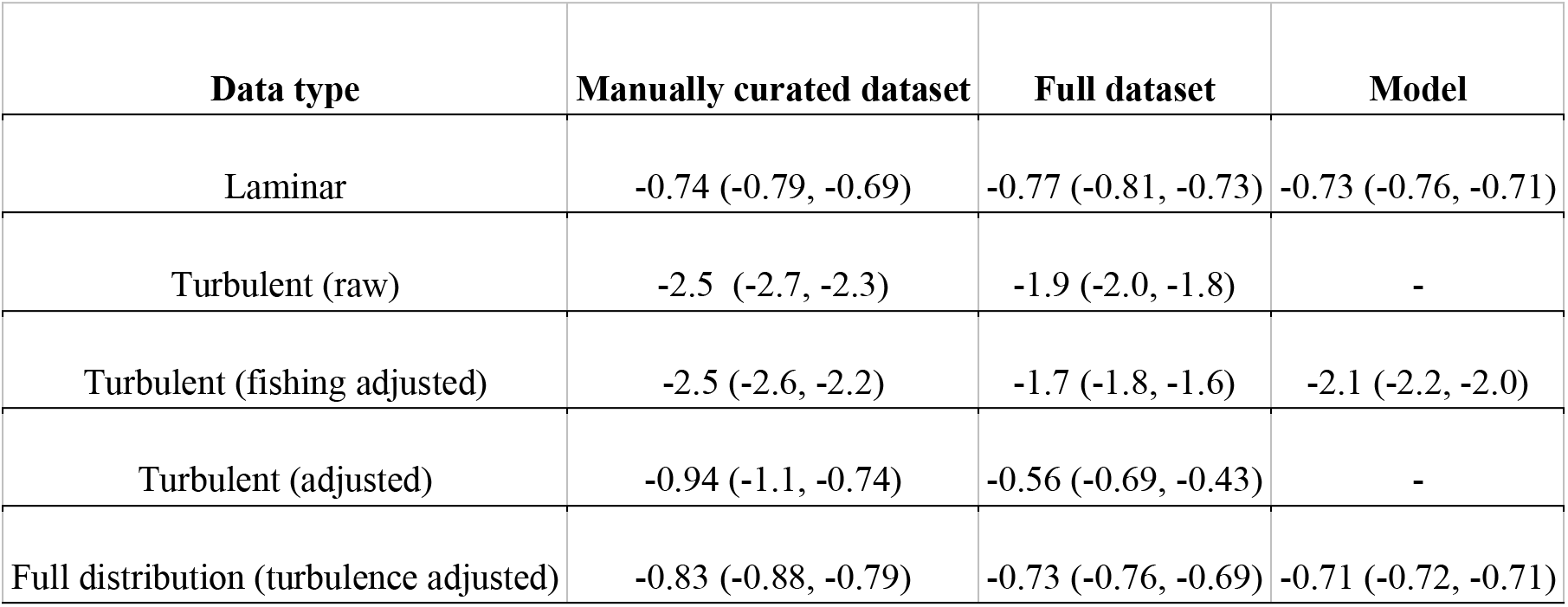
Estimates of the scaling exponent (*α*) with 95% confidence intervals for the empirical data (raw and adjusted) and the model simulated data, all calculated from 10,000 bootstrapped values (Methods).

| Data type | Manually curated dataset | Full dataset | Model |
| --- | --- | --- | --- |
| Laminar | -0.74 (-0.79, -0.69) | -0.77 (-0.81, -0.73) | -0.73 (-0.76, -0.71) |
| Turbulent (raw) | -2.5 (-2.7, -2.3) | -1.9 (-2.0, -1.8) | - |
| Turbulent (fishing adjusted) | -2.5 (-2.6, -2.2) | -1.7 (-1.8, -1.6) | -2.1 (-2.2, -2.0) |
| Turbulent (adjusted) | -0.94 (-1.1, -0.74) | -0.56 (-0.69, -0.43) | - |
| Full distribution (turbulence adjusted) | -0.83 (-0.88, -0.79) | -0.73 (-0.76, -0.69) | -0.71 (-0.72, -0.71) |

To find the cause of the break in the marine size-abundance relationship, we note that fishing has reduced the abundance of fish, pinnipeds, sea turtles and marine mammals by up to 99%^8^. We corrected for this by adjusting the abundances of impacted populations to pre-human impact estimates^8^. This caused a significant (*p* < 0.01) upward translation of the scaling line, removing the step break in the dataset and corroborating earlier findings^5^. However, whilst the translation is indicative of a decreased abundance of animals larger than 0.1 m, correcting for fishing did not result in a change in exponent, rather just a vertical shift in the data (Figure 1b, Table 1). The size-abundance distribution may be interpreted as an average or upper bound on local population densities^2^. The slope change is thus indicative of a constraint limiting nekton abundances which is not present in planktonic or terrestrial systems. To probe for a mechanistic explanation of the exponent change, we note that many aquatic organism scaling laws break at ≃ 0.1 m^6,9,10^; this size corresponds to the laminar-turbulent transition, where the change in the physical fluid environment causally affects the biology^6,10^. We subsequently tested the hypothesis that the change in scaling value is due to implicit and explicit costs associated with turbulence: that is, nekton must expend energy actively moving to match planktonic prey distributions, and that this expenditure propagates through higher trophic levels.

Aquatic predators and grazers are challenged by the chaotic nature of turbulence. As absolute abundances of resources scale similarly in three-dimensional aquatic and two-dimensional terrestrial environments^11^, their statistical distribution is scarcer in the three-dimensional ocean.

Plankton live within patches created by an interplay of physical and biological processes^12^. Within these resource hotspots, plankton foraging and movement is localised and constrained within the patch, allowing them to use hunting strategies such as chemotaxis or rheotaxis to maximise their food acquisition^13,14^; that is, plankton move passively with the turbulence that creates the aggregations. Beyond several millimetres and up to ten centimetres is a transition zone where eddies play an increasingly important role. Whilst they are below the swimming speeds of most fish, eddies on the scale of tens to hundreds of metres cause bulk transport and dispersal. Mesoscale eddies reach hundreds of kilometres in diameter and can move organisms hundreds or thousands of kilometres^15^. Food may not be transported, or it may be consumed and not replaced due to low light, low temperature or other unfavourable conditions^16^. Thus, nekton must migrate between patches to feed, which are continually and unpredictably dispersed, meaning they have resource encounter rates that typically cannot be bettered using local information^17^. Nekton live at a scale where the foraging landscape is highly fragmented and disordered due to these physical processes, and operate on biological timescales which are significantly longer than eddy lifespans^16,18^. As they are trophically linked to the plankton, they must actively work to overcome the dispersal, ultimately increasing their locomotory costs, which also grow with prey size^19^. Short distance dispersal within or just beyond local habitats is difficult to quantify. However, at a global scale, physical dispersal – and consequently the spatial distribution of plankton – follows the Kolmogorov power law for the turbulent energy cascade^12^. The overall effect is that dispersal, encoded here as the separation distance, is a key factor in nekton survival. We propose that resource acquisition forces nekton movement to follow the turbulence-driven distribution of plankton, increasing energy expenditure^20^, and consequently reducing available energy for growth and reproduction, which decreases abundances. The positioning of the break in the scaling relationship at the laminar-turbulent transition is consistent with this reasoning. Testing the hypothesis that turbulence increased the nekton slope by adjusting for the Kolmogorov power law, which affected small fish the least and large pelagics the most, removed the structural break in the distribution and resulted in a near-canonical exponent of *α* = −0.73 for the entire distribution (Figure 1d, Table 1).

To build a minimal model which captures this phenomenon, we note other scaling breaks for aquatic organisms^6^ also occur at 0.1 m due to movement changes at the laminar-turbulent transition^10^. The classical assumption that swimming is more energetically efficient than running^21^ does not consider drag, which increases with the square of velocity and carries extreme metabolic cost^22,23^. Research examining cost of swimming may also underestimate real-world metabolic effort for nekton as it frequently uses theoretically ‘optimal’ size-speed scaling^9^ rather than utilising empirical values which are steeper^6^. Finally, relative consumption rates are higher in oceanic than terrestrial environments, yet a steeper inverse scaling of nekton abundances in marine systems exists even at high resource densities^11^. This discrepancy has not been resolved but indicates there must be a significant energetic cost associated with living and feeding in oceanic environments that has not been considered. We incorporated classical formulations of swimming cost for organisms living in laminar and turbulent environments, together with foraging effort, into a size-dependent predator-prey model to assess these effects (Figure 2). In short, we expand the trophic transfer efficiency parameter, *ε*, in the classical Rosenzweig-MacArthur predator-prey model to account for energy diversion toward locomotion (Equation 1). In this equation, each parameter scales according to the length *l* of the organism (m), allowing it to be solved across the full size range.

**Figure 2:**
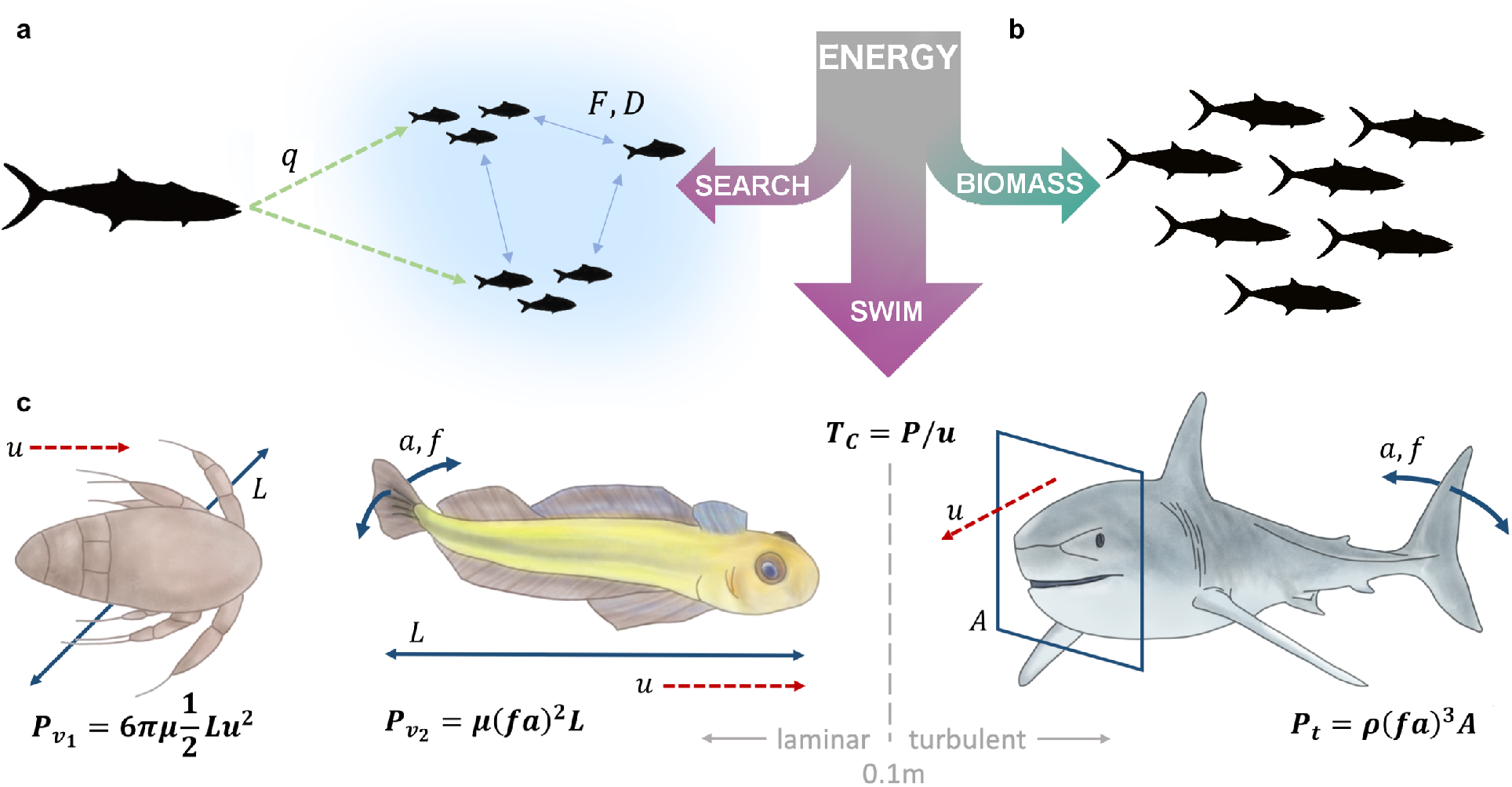
Energy partitioning - organisms have a finite energy budget which is split between movement and creation of new biomass. **(a)** A search effort term *q*(*F* – *D*) is described by the scaling of swimming speed (*q*), as well as parameters *F* and *D*, which denote resources’ fractal dimension (space-filling amount) and the physical dimension of the search space respectively. **(b)** Energy not spent on locomotion is utilised in reproduction and creation of new biomass. **(c)** Transport/swim cost (*T_c_*) is defined as power, *P*, divided by speed *u*. In the laminar regime, power for viscous paddlers, such as copepods, is described by length (diameter) *l*, speed, and viscosity *μ*. Viscous undulatory swimmer power (i.e. larvae or small fish) is given by kick frequency *f*, kick amplitude *a*, length, and viscosity *μ*. In the turbulent regime power is described by kick frequency and amplitude, frontal area *A* and fluid density *ρ*. We use these formulae to calculate size scaling exponents for swimming cost. The values can then be used in the master equation (Equation 1) to capture changes in energy partitioning across the size range.

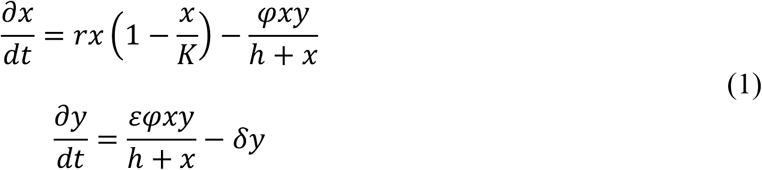

Capturing the shift in movement energy budgets from the laminar to the turbulent regime is achieved by using the relation *ε* ∝ *l^−c−q(F – D)^* (24). The exponent *c* accounts for the scaling of swimming cost relative to basal metabolic rate, and the term *q*(*F* – *D*) depicts resource search effort (Figure 2) (refer to Methods for the complete derivation).

Including locomotion cost for simulated predator-prey combinations from primary producers to blue whales reproduced the empirical results. Calculating the slope for model equilibria abundances in the turbulent regime resulted in a value of −2.1, consistent with the data (Figure 3a, Table 1). For the laminar model, and the turbulence-corrected predator-prey formulation across the entire data set, the slopes were −0.77 and −0.73 respectively, matching the empirical results (Figure 3b, Table 1). In our model, living in a turbulent fluid regime impacts the system by translating the predator abundances downward. This means prey support fewer predators in a turbulent environment than they would in viscous or terrestrial regimes because of the increased energetic costs of foraging in turbulence. Increasing locomotion energy budgets decreases biomass transfer to higher trophic levels where reduced prey availability places even more restrictions on energetic resources^19^, pushing large marine organism abundances closer to an unviable population threshold where natural population fluctuations also render them more vulnerable to extinction^25^.

**Figure 3:**
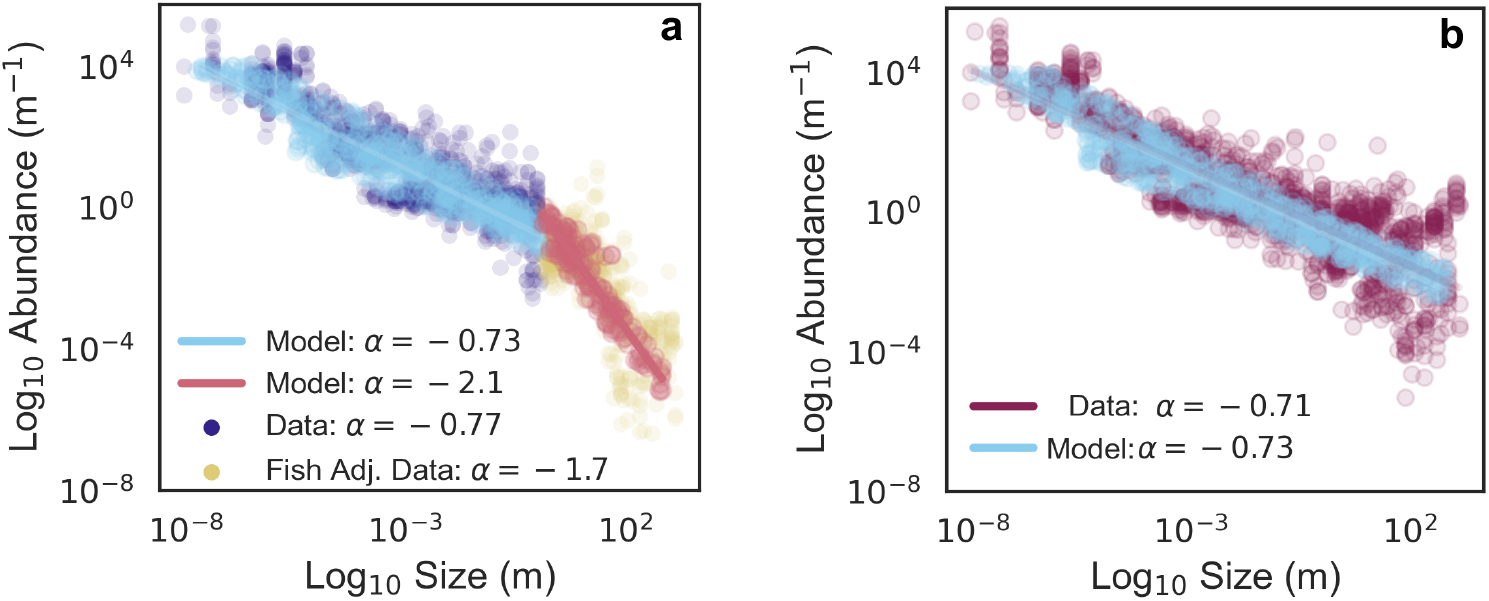
Rosenzweig-MacArthur model results. **(a)** Plankton (dark blue) with fishing corrected empirical data (yellow), the laminar model simulated data (*l* < 0.1 m, pale blue) and turbulent model simulated data (*l* ≥ 0.1 m, red). **(b)** Fishing and turbulence corrected data (purple circles), are shown with the model simulated data (pale blue) superimposed over the data-fitted regression line.

As our model includes a parameter for resource density, direct impacts of overfishing may also be incorporated. We find that whilst heavy fishing could theoretically perturb the size-abundance scaling value by decreasing resource saturation *F*, the search effort multiplier *q* is ~0.17 (relative to mass). This means it is a slow parameter, which also reaches an asymptotic value as *F* → 0. Hence, whilst fishing removes biomass, our integrated model indicates it could only perturb the size-abundance scaling law by ≈ −0.2 before the asymptote is reached. This is an order of magnitude less impact than turbulence effects, and entirely consistent with what we observe with our data (Table 1).

A complicating factor with our analysis is that organisms and biomes are not fixed physical or chemical variables. Their characteristics can change in response to environmental pressures. Ecosystem-wide size shifts in size-abundance relationships may be exacerbated by compensatory genetic changes, particularly when they have occurred under strong selection pressures such as fishing. Such a fisheries-induced evolution (FIE) causes further size reduction and earlier maturation age^26^, which could alter the scaling relationship. To assess the relative impact of FIE, we extracted data from 113 time series for 10 commercially exploited species of fish, and assessed global changes in size and age at maturation. There was a mean decline of 11% in size or age at maturity, when accounting for gender, species, and length of study (Methods). The results from 10 of the 14 studies led to the conclusion that these changes were attributable to fishing pressure^26^. In considering FIE’s contribution to universal size-abundance scaling, the breadth and size of our dataset gives insight into the signal-to-noise ratio for this problem. It would be extremely challenging to detect shifts in a global scaling law over the restricted size range of 0.1 to 2 m used for FIE impacts. While prior research suggests that FIE can perturb local scaling properties^27^, we argue an 11% impact (or even significantly greater) would not be enough to shift the global size-abundance scaling value of nekton by −1 or more. We conclude that scaling alterations occurring due to FIE would be small relative to the turbulence effect explored in this paper.

Global size-abundance laws provide a different form of ecological insight to that given by local scaling behaviour, as they capture macroscale, aggregate processes rather than examining small-scale drivers such as inter- and intra-specific trait variation^2^. In this context, we introduce turbulence, and its impact on energy and movement cost for large organisms, as a novel but important process to consider for ocean ecosystems. Climate change impacts have the potential to exacerbate these costs, as current and predicted increases in ocean surface energy^28^ will increase nekton foraging and locomotion costs^29^, whilst warming temperatures increase respiration rates, reduce global primary productivity^30^, and cause greater resource patchiness^31^, forcing increased movement cost. Turbulence may thus reduce the capacity of nekton to withstand fishing pressure as we begin to observe oceanic anthropogenic impacts classically associated with terrestrial systems, including loss of large apex predators, shifts to smaller size, and a faster onset of sexual maturity. We propose that a deeper understanding of the role physical mechanistic processes play in structuring marine ecosystems will be necessary when formulating strategies to preserve biodiversity and retain the productivity of ocean resources in future.

## Acknowledgements

Stephen J Hall, Peter G Strutton, Susan D Willmott, Xiaoke Hu, Song Sun, and Lisa Dann made helpful comments on early versions of the manuscript. This work was supported by Flinders University, CNRS and Australian Research Council grants to JGM, LS and JRS. Matthew D Russell, Shareena J White, Tomoyo Segawa and Rachel J Pillar helped with data collection.

## Author contributions

JGM, JCM, JRS & LS conceived the work

JCM, JGM & JRS wrote the paper

SRG, TJL, PJR, TCJ, JSP, BR, CH, KN, VvDV, NPC, KMW, RJS, CLB, EP, JRS and CMC gathered the data and helped with the analysis

JCM developed the model, gathered data, and did the analysis

MK contributed to model development

MT helped with analysis and contributed to writing, interpretation and insight

LS helped with analysis, and contributed to writing and interpretation and insight

## Data and materials availability

All data is available in the supplementary materials or online. Code is available at https://github.com/jcmckerral.

## Methods

The reader may refer to Extended Data, Table S4 for symbol definitions used throughout the Methods. All statistical testing was conducted in MATLAB R2019b (Mathworks), and code is available at https://github.com/jcmckerral.

### Data

#### Data sourcing and aggregation

To assess the size-abundance scaling relationship, we examined data for over 2100 species, encompassing over 800 genera (bacteria/viruses excluded from diversity counts) (Extended Data, Table S1, S7). For quality purposes, we undertook analysis with two datasets. The first was manually curated from over 200 articles to ensure there was not systematic bias within database sources, and consists of 1719 size-abundance pairs across 700+ species (Extended Data, Table S1). The second dataset expands on the first via the inclusion of a further 13,455 entries predominantly sourced from online databases, for a total of 15,174 data points (Extended Data, Table S7). Five databases were used: IMOS (flow cytometry and zooplankton)^32,33^, Tara Oceans (flow cytometry)^34^, Phytobase^35^ for phytoplankton, a global diatom database^36^, and a reef fish dataset^37^. Size data was taken from the same source as the abundance data, or if it was not included, we assigned the average adult size for that taxon referenced from WoRMS^38^, fishbase^39^, or (36) for diatoms. All database entries which dated pre-2000 were removed to reduce the chance of methodological/quality control problems being introduced from older data. For Phytobase entries, any data with the flags ‘unrealistic day or year’ and ‘presumably sedimentary’ were deleted; we note this particular database is otherwise well suited to this application as capturing local diversity patterns is not critical for global size density analyses^2^. For the flow cytometry data, any entries which had not undergone or passed quality control checks were removed.

Next, we outline pooling information for taxonomic/sampling groups. For most nekton, abundance estimates were given at the species level, with the exception of hard-to-differentiate taxa, e.g. striped/common dolphins. Unless the data had been provided that way by the primary source, no averaging or grouping was undertaken. For bacterial and viral abundances, we elected to use flow cytometric data rather than DNA-based methods, as the high variance in copy numbers of marker genes in prokaryotes precludes reliable estimates. (Note that size measurements for bacteria and viruses were given by microscopy-based sources, not flow cytometry.) In addition, defining ‘species’ grouping is inherently problematic for microbes. No manually curated data was aggregated unless that was its original format. For the databases, we pooled according to the following principles. Firstly, we took taxa abundance averages by year and location. A single location was taken to be one station, or the same degree of latitude/longitude. We averaged at the lowest available taxonomic level (usually genus for organisms <5E-4m, and species for anything larger), and selected taxa which, together, provided >90% of the total abundance of that sample to avoid skewing with singletons; this also aligns with the principle of size-abundance distributions often being representative of abundance average or upper bounds^2^. The exceptions to this pooling rule were for targeted flow cytometry counts of abundant cyanobacteria (*Prochlorococcus*, *Synechococcus*) which we included as is.

Abundance data is localised, hence spatial and temporal variation across local snapshots captures natural variability of populations across space and time. Therefore, the inclusion of data from different environments, e.g. tropical/temperate, or low/high biomass regions, or across different sampling efforts, is suitable – and even desirable – as the goal is to build the universal distribution, which should ideally contain a broad spread of data^2^. Given the similarity between the manually curated and complete database results, and the generally well-behaved nature of the model statistics (Figures S1-S3), we elected not to transform or apply other corrections to the data. We acknowledge there is certainly variance introduced from species trait differences, and potentially from inconsistencies from underlying experimental methods. However, these impacts would remain with noise factor of this dataset. Furthermore, whilst more targeted studies can be sensitive to this variance due to scaling size range and data limitations (e.g. bony fish, at ~3 orders of mass magnitude)^2^, fitting the scaling exponent over 23 orders magnitude, with this quantity of aggregated data, drastically mitigates the effect of any one source of error. Notably, the noise was sufficiently low for a strong statistical signal without the need for any manipulation, which could introduce other errors or biases, and reduce transparency of the result.

#### Standardisation and units

Due to the large mass range (> 23 orders of magnitude), measuring uncertainty in the body mass of microorganisms^40^, and to ensure units were consistent in downstream analyses, we used body length, *l* (m), as the measure of organism size. To accurately compare data sets where abundance measurements were presented either as species numbers per unit volume or per unit area, and to account for organism behaviour, we calculated the separation distance, *d* (m), between organisms as a proxy measurement for abundance. To calculate separation distances, it was assumed the spatial distribution of organisms followed a Poisson distribution. Thus, the separation distance for organisms where abundance was measured per unit area was given by *d* = *C*^−1/2^, and per unit volume, *d* = *C*^−1/3^. Under the assumption that organism mass is approximately proportional to organism volume, the transformation of both axes in the size-abundance plot is the same. Therefore, our standardisation to length does not change the empirical scaling values, nor does it disproportionately impact one part of the distribution, but instead ensures consistency with units in the physics-based processes and derivations used in the analyses. We acknowledge that organism mass and length generally do not have a perfect cube root relationship. However, this is a standard transformation utilised when investigating bioenergetics of swimming organisms^9^; we also note that any deviation from a cube root relationship would be applicable across the full distribution and therefore not change the key outcomes of our analysis relating to the structural break.

We now discuss the raw data and the potential errors that may have arisen due to this standardisation. Plankton data was near universally presented by volume; we note that plankton distributions are by definition patchy and this variance far exceeds that of methodological error. Volume-based measurements in the reef fish dataset were based on study areas <30m deep and already undergone significant quality controls for accuracy; we did not undertake any further corrections. We assumed volume-based data for small nekton in the manually curated literature data did not require further adjustments. We acknowledge some small amount of error may have been introduced under this assumption in the event that depths were incorrectly measured, but note that (a) in the context of incorrectly measured depths, the cube root transformation reduces the impact of that error and (b) the data covers approximately 0.5 of an order of (length) magnitude, meaning that impacts on the full distribution would be minimal, particularly after log-transformation. For marine megafauna, only studies using standard methodologies according to transect/aerial surveys were included. It is to be noted that most of the length- or area-based abundance measurements in the dataset were aerial survey data of marine mammals, and not of benthic organisms.

#### Power law model fitting methods

To determine the scaling relationship across the dataset, organism length was plotted against the inverse of the separation distance 1/*d* (m^−1^) on a logarithmic scale, so that *d* ∝ *l^−τ^*, where *τ* is a scaling exponent. Note that we consider a global, bivariate, size-abundance distribution more commonly applied in terrestrial settings, and not the univariate size distribution often studied in aquatic environments^2^. As the data is bivariate, the methods developed for univariate distribution fits are not directly applicable^41^. Regression methods are standard for the bivariate case, and may be used provided the dependent variable contains higher measurement error than the independent variable^42^. Therefore, following a residuals analysis, the models for plankton and nekton were fitted using ordinary least squares on the log-transformed data (residuals plots provided in Figures S1-S2). For the fits, a balanced subsampling routine was used to ensure an even spread of data across the distribution and improve fit quality^43^. We did not use a naive with-replacement bootstrapping routine as this would simply bias the sampling towards whichever data (taxa and/or sizes) were most frequent in our database. Furthermore, as large databases typically had large groups of data clustered together (e.g. Figure 1a, where various clumps of data may be observed), subsampling mitigated against one database, taxon, or size class dominating the fit. The data was stratified by organism sizes, and by taxa. We then randomly sampled *m* data points (without replacement) such that the quantity of data per (log)bin was uniform across the full size range and balanced the probabilities of sampling from different taxonomic groups. The optimal subsampling size *m* is denoted by *m* = *kn^κ^*, where *n* is the size of the dataset being drawn from, *k* = 3, and *κ* = 0.5 (43, 44). We then generated 10,000 parameter estimates for each model, where each estimate was created from subsampled data, for the laminar regime *l* < 0.1, turbulent regime *l* ≥ 0.1, or complete size range. Percentile confidence intervals (95%) were created from the bootstrapped statistics. Representative linear model statistics are available in Tables S5-S6, and bootstrap histograms in Figure S3. For the *α*-estimates from the Rosenzweig-Macarthur simulated data, we randomly generated *m* datapoints (matching the empirical subsample sizes) for the laminar, turbulent, and full size ranges. Confidence intervals were generated from fitted linear models on 10,000 model runs for each *α*-estimate.

#### Structural break

We used MATLAB’s fminbnd function to find the segmented regression breakpoint which minimised MSE. This was bootstrapped for a percentile-based confidence interval on log-transformed, subsampled data (sampling method as for regressions).

### Correction for Fishing

To investigate the impact of fishing on the observed scaling relationships, organisms were assigned to groups of impacted large marine animals according to standard conventions^8^. These included organisms such as fish, sharks, pinnipeds, whales, sea turtles and sea birds. Separation distances were corrected for each group to reflect theoretical historical abundance values, assuming losses ranging between 50 and 99.7%^8,45^. Where no specific loss estimate was available, the mean decline for all large marine species (89%) was allocated^8^.

### Correction for Aquatic Turbulence

The influence of aquatic turbulence on the scaling relationship for nekton was addressed by applying a phenomenological correction for the −5/3 relationship arising from the Kolmogorov power-law of the inertial subrange of the energy spectrum^46^. The spectral energy density, a proxy of the variance of the variable under consideration, i.e. turbulent velocity fluctuations in the framework of fully developed turbulence, is given by *E*(*k*) = *C_k_ε*^2/3^*k*^−5/3^, where *C_k_* is the Kolmogorov constant (~1.5), *ε* is the turbulent kinetic energy dissipation rate and *k* is the wave-number (2π/eddy diameter, rad.m^−1^)^46,47^. Here we approximate this relationship as *E*(*k*) ∝ *k*^−5/3^, providing a dimension of *m*^−1^(47). The spatial distribution of plankton has been observed to follow the same power law^12,48^, and the separation distance *d* as a function of size (both units in m) may therefore be considered as an implicit measure of the effect of dispersion due to turbulence. Thus, by considering *d* ∝ *k*^−5/3^ we undertook a phenomenological correction for the abundances of nekton, whose foraging effort is impacted by the turbulent dispersal of plankton, by subtracting the Kolmogorov power law, intersecting at *l* = 0.1 m, and calculated an adjusted scaling value for the entire data range.

### Rosenzweig-MacArthur model

We used the classical Rosenzweig-MacArthur model to investigate the effect of turbulence on population dynamics and size-abundance relationships for consumer and resource pairs, from phytoplankton to whales. This formulation allows us to use previously defined allometric laws to generate a global size-abundance distribution. Despite the number of assumptions inherent in allometry, we note that macro-scale models parameterised by size have been found to outperform those which are defined based on species-specific traits and are also significantly more parsimonious^49^. To maintain consistency in units across empirical data, model, and adjustments, size was given by (standardised) length in m and abundance was defined as organisms per meter (*n*. m^−1^), i.e. the inverse of separation distance, rather than mass (kg) and biomass (density, kg. m^−3^). The base ordinary differential equation contains strictly positive parameters and is described by:

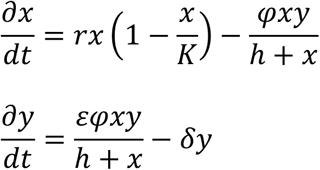

where *x* and *y* are resource and consumer (predator) abundances, respectively. The parameter *h* denotes the half saturation, whereas *K* is the carrying capacity, *r* and *δ* are birth and death rates, *ε* the conversion efficiency, and *φ* the maximal consumption rate.

Each of the parameters follows scaling models according to the size (*l*, m) of the resource (*l_x_*) or consumer (*l_y_*), such that *i* = *i_o_l^σi^*, for some parameter *i*, coefficient *i_0_* and exponent σ_i_. Scaling properties can change according to factors such as primary production rates, temperature, habitat complexity, among many others^3^. A constant temperature was assumed, and resource-consumer size ratios between 0.01 and 0.5 (corresponding to prey-predator mass ratios of 1E-6 and 0.1 respectively), as scaling laws can change when the predator is smaller than the prey. Exponents were given by representative values from previous research, which was typically specialised on deriving empirical scaling for that specific parameter (Extended Data, Table S2). As our dataset ranges over more than 23 orders of mass magnitude, where there was some variability across literature scaling models, our study used the exponent values which were most consistent across the size range. Values chosen were (i) frequently reported with consensus (*r*, *δ*, *φ_v_*, *K*), (ii) midrange (*h*) or (iii) specifically calculated for aquatic vertebrates (*φ_t_*). Noting that rate-related parameters (*r*, *δ*, *φ_v,t_*) will scale faster with length than mass, the scaling values are given as follows: 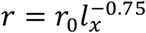; 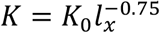; 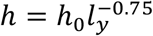; 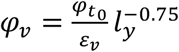; 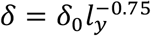; 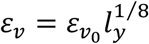. (Length scaling values of *-3/4* are equivalent to mass scaling values of −1/4.) We assumed carrying capacity scales according to −3/4 as per null metabolic expectation, but note here that it does not impact equilibria values in the Rosenzweig-Macarthur system of ODEs (although it does affect behaviour of the limit cycle). The scaling values for two parameters change between the viscous and turbulent regime (organism length >0.1 m): 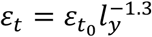 and 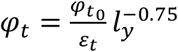. Please refer to Extended Data, Table S2 for literature sources for exponents, and the biophysics section below for the derivations of *ε* scaling exponents. Under this parameterisation, there is a switch to a positive maximal consumption rate (*φ*) in the turbulent regime. This has been previously noted in the functional response literature. Whilst invertebrates and microorganisms typically scale with a −0.75 (length) exponent, which matches null model predictions derived under metabolic theory, data for macroscopic fauna in aquatic environments display positive scaling; our derived *≈* 0.55 exponent falls within observed empirical ranges^11,49,50^. Refer to Extended Data, Table S2 for more information.

The parameter scaling coefficients were standardised against phytoplankton/zooplankton models to ensure the boundary value for primary producers was feasible. The smallest primary producer (i.e. 0.7 – *1 μm* in length) was assumed to be the cyanobacterium *Prochlorococcus*^51^. For coefficients, biomass was divided by species mass to obtain the number of organisms. Model equilibria were calculated using analytical formulae solutions.

### Locomotion cost: biophysics derivations for the model

To derive the biophysics portion of the model we integrate models across several disciplines. We use scaling of mass throughout this section to remain consistent with the literature, unless otherwise specified. To account for movement cost in the Rosenzweig-MacArthur system, we consider locomotion energy budgets across the whole size range (bacteria to whales). If movement energy usage scales equivalently to basal metabolic processes, its impacts would not be noticeable. However, if it scales differently, some of the energy previously used to create new biomass would instead be diverted to locomotion. Alternately, if locomotion were to become more efficient, additional energy could be provided for biomass. This can be seen by examining the gross metabolic power of an organism:

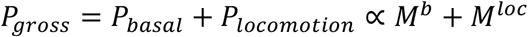

Normalising by *P_basal_* results in:

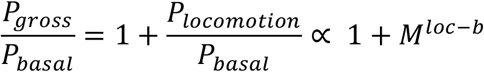

If there is a discrepancy between the power exponents, the (relative) locomotory power consumption will change across the size distribution.

This deviation can be captured within the parameter for biomass transfer efficiency ε. To achieve this, we use a classical ecological relation, which links basal and locomotory metabolic cost to abundance^24,52^:

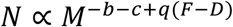

In this master equation, *N* is the population abundance, and *c* is the relative transport cost scaling. We have *c*:= *p* – *b*, where *b* is basal metabolic scaling, and *p* is the scaling of transport cost (*T_c_*) defined below. The term *q*(*F* – *D*) describes search effort, including *q*, swim speed scaling, and the parameters *F* and *D*, which describe density/fragmentation and dimensionality of the resource space. Note that if the term -*c*+*q*(*F* – *D*) equates to zero, classical population dynamics apply. That is, the standard Rosenzweig-MacArthur system, with a typical value of *ε* e.g. the prey-predator size ratio. However, when it is non-zero, it captures the shift in locomotion energy allocation across the size distribution. This provides the following relationship for *ε*:

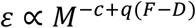

In the subsequent derivations for the exponents of ε, we use empirical swim speed scaling results from Andersen *et al*.’s (2016) review of marine scaling laws: 1/4 and 1/6 for viscous and inertial swimmers respectively. This is important because it suggests the scaling of real-world nekton swimming speed is steeper than what would be theoretically derived for maximum efficiency. ‘Optimal’ speed scaling would be given as 5/24 and 1/12 for viscous and inertial regimes (calculated according to methods in Bale *et al*. 2014 Supplementary Information, under the assumption of a 3/4 basal metabolic law).

#### Search effort scaling (q(F – D))

The parameter *q* is the scaling of swimming speed. The dimensionality of the space, *D*, is taken as 3 for the turbulent regime. In the laminar/viscous regime, we consider *D* = *D’* = 2.4, to account for the patch constraint and the fact that organisms can use local information to optimise their hunting strategies^24^. We set the fractal dimension of the space, *F*, to a mid-range value of 1.9^24^.

#### Transport cost scaling (p, c)

In this section, *μ* and *ρ* denote the viscosity and density of the liquid respectively. For the purposes of this study, we assume all physical fluid properties are constant as changes in transport cost due to pressure, salinity, or temperature fluctuations at depth or in tropical versus polar regions are negligible relative to the effect of changes in size of the organism.

Transport cost is defined as *T_c_* = *P/u* where *P* is power and *u* is swimming speed^9^. The master equations for the power of swimmers in the viscous regime are given by *P_v.und_* = *μ*(*fa*)^2^*l* for undulatory swimmers^9^ and 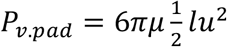 for paddlers^53^. Here, *f*, *a* and *l* are the kick frequency, kick amplitude and body length respectively. By using the classical relationship determined by Bainbridge^10,54^

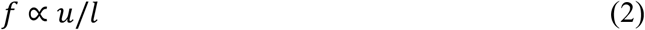

and assuming changes in the length measurements *a*, *l* are scaling approximately proportional to *M^1/3^*, we have *P_v.und_ ∝ M^5/6^* and *P_v.pad_ ∝ M^5/6^* after value substitutions. That is, the power cost scales equivalently for paddlers and undulatory swimmers in the viscous regime.

For the turbulent regime, the power of inertial swimmers is given by *P_t_ = *ρ*(*fa*)^3^A*, where *A* is the frontal area of the organism (scaling as *M^2/3^* accordingly)^9,10^. Once again, we use Equation 2 and substitute in mass scaling values to obtain *P_t_ ∝ M^7/6^*. Using the definition of transport cost, we obtain *T_c_v__* ∝ M^7/12^ for organisms in the viscous environment and *T_c_t__* ∝ *M* for the turbulent environment. As the units for *T_c_* are J/m, it is possible to make it unitless via multiplying by 1/*ρν^2^*, which is a constant under our assumptions of fluid properties.

This means that: 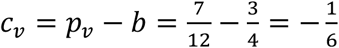, and 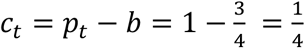.

With the values for *c*, *q*, *F* and *D*, we can now calculate the scaling for *ε* in the viscous and turbulent regime, which we then convert to length scaling:

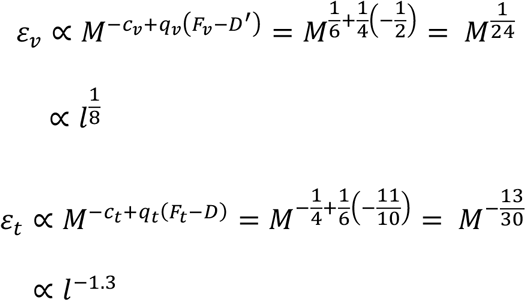

We switch between the parameterisations at the length of 0.1m, corresponding to the transition from laminar/mixed fluid regime to a fully turbulent flow of *Re* > 1000. Finally, normalising constants *ε_t_0__*, *ε_v_0__* set initial values. The resultant mean, maximum and minimum conversion efficiencies are 0.09, 0.2 and 4E-3 respectively, which are within expected literature values^55^.

### Fishing-induced evolution

Fishing-induced evolution (FIE), specifically, quantifying phenotypic change, was assessed by extracting size/age at maturity data from 113 time series taken from 15 studies (Extended Data, Table S3). In some cases, this was provided as probability norms of weight or length at 50% maturity (Wp50 or Lp50). Time-series with large gaps or fewer than 20 measured time points were excluded. Data was manually extracted using WebPlotDigitizer (v 3.12) and visually verified by replotting and super-positioning over the original. For plots without discrete data points (i.e. smooth line graphs), one data point per year was used. Each time series was normalised and then split in two halves, for which mean values were calculated for the first/second half of study period. This was imported into a data structure consisting of the mean values, data type (size or age at maturity, 50% maturity, Wp50 or Lp50), gender, species, and length of study. For testing the difference in means between the first and second halves of a study period, data was firstly assessed for normality by using a 2-sided Kolmogorov-Smirnov test (n=113, critical value=0.1262, observed values 0.0774 and 0.0958 for pre- and post-respectively, MATLAB R2016b, Mathworks). A paired t-test (SPSS 24.0.0.0, 2017) indicated a 10.6% shift in mean value in the second half of the study period (df=112, 95% CI (9.4,11.9), 2-tailed, t-statistic −17.374, p<0.001).

